# Femora from an exceptionally large population of coeval ornithomimosaurs yield evidence of sexual dimorphism in extinct theropod dinosaurs

**DOI:** 10.1101/2022.09.20.508522

**Authors:** R. Pintore, R. Cornette, A. Houssaye, R. Allain

## Abstract

Sexual dimorphism is challenging to detect among fossils, due to a lack of statistical representativeness. The Angeac-Charente *Lagerstätte* (France) represents a remarkable “snapshot” from a Berriasian (Early Cretaceous) ecosystem and offers a unique opportunity to study intraspecific variation among a herd of at least 61 coeval ornithomimosaurs. Herein, we investigated the hindlimb variation across the best-preserved specimens from the herd through 3D Geometric Morphometrics and Gaussian Mixture Modelling. Our results based on complete and fragmented femora evidenced a dimorphism characterized by variations in the shaft curvature and the distal epiphysis width. Since the same features vary between sexes among modern avian dinosaurs, crocodilians, and more distant amniotes, we attributed this bimodal variation to sexual dimorphism based on the extant phylogenetic bracketing approach. Documenting sexual dimorphism in fossil dinosaurs allows a better characterization and accounting of intraspecific variations, which is particularly relevant to address ongoing taxonomical and ecological questions relative to dinosaur evolution.

## Introduction

Dimorphism has been reported in every major dinosaur clade and has often been attributed to sex-specific variation (Dodson, 1976; Chapman et al., 1997; Bunce et al., 2003; Padian and Horner, 2011; Knell and Sampson, 2011; Knell et al., 2013; Mallon, 2017; Saitta et al., 2020). However, recent studies have demonstrated that most of the documented cases of sexual dimorphism in extinct dinosaurs were most likely biased by ontogenetic changes, taphonomic deformations and small sample sizes, which substantially affect the representativeness of the inter- and intraspecific diversity, and undermine statistical analyses (Griffin and Nesbitt, 2016; Hone and Mallon, 2017; Saitta et al., 2020). For example, a discrete and binary variation between gracile and robust morphologies of bone scars, mostly at the level of the lesser trochanter, has frequently been inferred, with more or less confidence, as sexual dimorphism in various ceratosaurian theropods and non-dinosaurian dinosauriforms (Colbert, 1990; Raath et al., 1990; Benton et al., 2000; Britt et al., 2000; Carrano et al., 2002; Piechowski et al., 2014). More recently, Griffin & Nesbitt (2016) demonstrated that this feature no longer appeared dimorphic when accounting for ontogenetic series in the silesaurid *Asilisaurus*. At a larger scale, Mallon (2017) performed a statistical investigation on a large set of studies that hypothesized sexual dimorphism based on a wide diversity of anatomical proxies across the major clades of non-avian dinosaurs. However, among the 48 described occurrences, only nine datasets were suitable for statistical test, among which only one was considered to rigorously demonstrate dimorphism. Indeed, the combination of a principal component analysis and a mixture modelling analysis highlighted that the shift in posterior inclination between the 8^th^ and 9^th^ dermal plates of *Stegosaurus mjosi* was best explained by a bimodal distribution. Yet, there is not robust evidence to postulate that the dimorphism shown in dermal plates would be sex-specific (Saitta, 2015). As a consequence, it appears that no dataset enabled to rigorously demonstrate the presence of sexual dimorphism in non-avian dinosaurs (Hone et al., 2020). According to Mallon (2017), one should review three issues when demonstrating sexual dimorphism on extinct organisms: 1) sample size in order to ensure population representativeness; 2) methodology in order to use only suitable analyses to study sexual dimorphism, such as mixture modelling; (3) any other intraspecific morphological variation such as ontogeny and pathology, as well as taphonomy.

Here, we studied the intraspecific femoral variability among a remarkable population of ornithomimosaurs (Allain et al., 2022, 2014) from the Angeac-Charente *Lagerstätte* (Lower Cretaceous of France). Rozada et al. (2021, 2014) demonstrated that at least 61 ornithomimosaur individuals belonged to the same herd and were deposited in a mass mortality event relying on several evidences (e.g., very limited transport; quality of bone preservation; abundance of individuals with a high skeletal representation preserved in a restricted spatial distribution; catastrophic age profile of the group; deposition of sediment and bones under coeval; poorly oxygenated burial and diagenesis conditions given by their rare earth elements and Yttrium profiles). Thus, the ornithomimosaur herd of Angeac-Charente represents a unique occasion to study subtle parameters such as intraspecific variability in extinct dinosaurs. Moreover, the exceptionally high minimal number of individuals among the herd offers a singular opportunity to test for the presence of dimorphism and characterize its variation.

We used a 3D Geometric Morphometric (3D GMM) approach that combines anatomical landmarks and sliding semilandmarks along curves and surfaces on both complete and fragmented femora and tibiae (Fig. S1A-B) (Gunz et al., 2005; Gunz and Mitteroecker, 2013). This method is well suited to study biological objects, including limb bones, and to detect subtle intraspecific shape variations (Zelditch et al., 2012; Botton-Divet et al., 2016) such as dimorphism (Fabre et al., 2014). We then investigated the resulting dataset using Principal Component Analyses (PCA) and Gaussian Mixture (GM) modelling as clustering analyses. This clustering analysis calculates the number of Gaussian distributions present in a dataset by maximum likelihood estimations and has been demonstrated as a well-suited method for the identification of dimorphism (Godfrey et al., 1993; Dong, 1997; Fabre et al., 2014; Manin et al., 2016; Mallon, 2017; Saitta et al., 2020)

**Institutional abbreviation**: ANG: Angeac-Charente Collection, Musée d’Angoulême, Angoulême, FR

## Results

We highlight a dimorphic variation in femora from the ornithomimosaur herd of Angeac-Charente (Fig. 1A-B). This dimorphic variation is localized along the diaphysis (i.e., lateromedial curvature) and toward the distal epiphysis (i.e., lateromedial width) of the femur (Fig. 1C-D). Distributions along the PC1 of complete femora (28.8%) and distal epiphyses (27.9%) are best described by two clusters with a ratio close to 1:1 according to mixture modelling analyses (see Table S1 for details). PC1 scores from both analyses are not significantly correlated to the log centroid size, indicating that size-related effects have no impact on the observed dimorphism (*p-value* > 0.1 for complete femora and distal epiphyses, Table S1).

**Figure 1:**
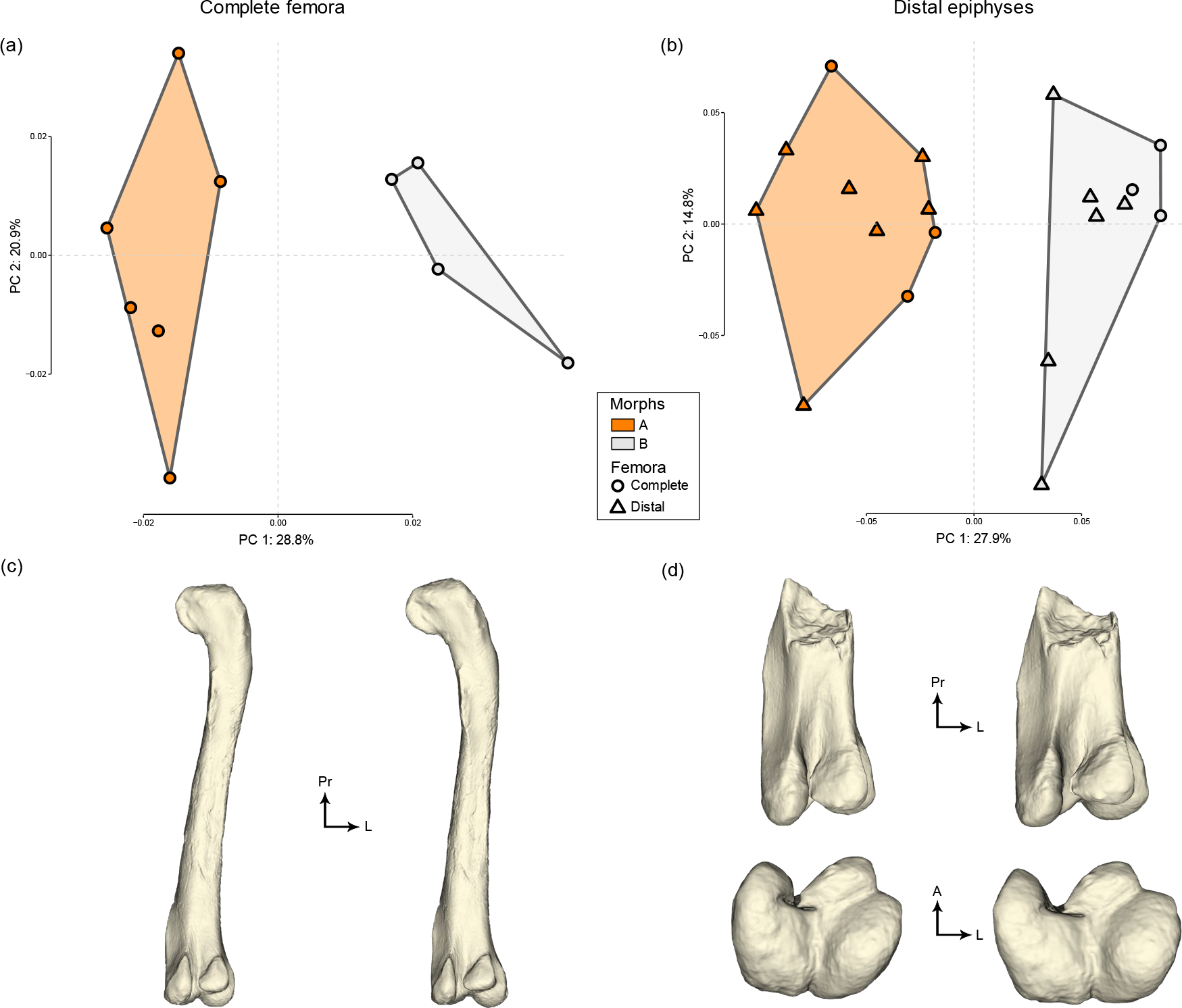
The two first axes of the PCA for A) complete femora and B) distal epiphyses; Minimal (left) and maximal (right) mean shapes per group for C) complete femora in posterior view and D) distal epiphyses in posterior (top) and distal (bottom) views. Abbreviations: L, lateral; P, Posterior; Pr, proximal.

The most important morphological variation of complete femora is a medial to lateral curvature of the femur (Fig. 1C). The proximal third of the femur appears deviated toward the lateral side in specimens on the negative part of the axis, whereas specimens located on the positive part have straight to medially curved femora (Fig. 1C). Coincidentally, the femoral head is directed medially in the negative cluster while it is inclined ventromedially in the positive one (Fig. 1C). Regarding distal epiphyses, we selected six (out of 10) epiphyses from complete femora because the other four were taphonomically altered or pyrite encrusted only in the distal area, which would appear relatively more important in analyses restricted only to this area rather than on the complete morphology (Table S2). Nevertheless, for distal epiphyses the most important morphological variation along PC1 is the expansion of the lateromedial width relative to the anteroposterior length, which is greater in specimens on the positive part of the PC1 axis than on the negative one (Fig. 1D). In addition, we highlight that the six distal epiphyses from complete femora are consistently attributed to the same clusters between the two analyses (Fig. 1A-B; Table S2). Hence, our study shows that the straighter the shaft is, the more robust the epiphysis is, and that this relationship is dimorphic.

However, there is no robust bimodal distribution on proximal epiphyses, as shown by the GM analyses (Fig. S2; no consistency in the specimen attribution between complete femora and proximal epiphyses). Similarly, there is no dimorphism in the morphological variation of complete tibiae (Fig. S3) along PC1 (24.1%) and PC2 (20.0%).

## Discussion

The closest extant relatives of non-avian dinosaurs are known to display sexual dimorphism with more or less visibility: birds display variation in their plumage and skeleton (Schnell et al., 1985; Owens and Hartley, 1998; Dunn et al., 2001; Székely et al., 2007; Clarke, 2013; Duggan et al., 2015; Manin et al., 2016; Hone and Mallon, 2017; Elzanowski and Louchart, 2022), whereas the variation is restricted to skeleton in crocodilians (Fitch, 1981; Farlow et al., 2005; Cox et al., 2007; Prieto-Marquez et al., 2007; Bonnan et al., 2008; Hone and Mallon, 2017; Hone et al., 2020). The extant phylogenetic bracket (EPB) of non-avian dinosaurs (Witmer and Thomason, 1995) thus implies they were sexually dimorphic too (Hone and Mallon, 2017; Hone et al., 2020).

A femoral dimorphism of the same nature was demonstrated to be sex-specific among populations of extant tetrapods such as carnivorans and primates. Dimorphism in the femoral obliquity (also termed “bicondylar angle”) was observed in humans, for which females had higher angles than males (Parsons, 1914; Tardieu et al., 2006; Hunt et al., 2021). Moreover, a higher lateromedial width of the distal epiphysis (also termed “epicondylar width” or “bicondylar breadth”) was demonstrated to vary between sexes in grey wolves and other carnivorans, as well as in primates (Alunni-Perret et al., 2008; Gaikwad and Nikam, 2014; Morris and Brandt, 2014; Cavaignac et al., 2016; Morris and Carrier, 2016). Whereas no similar sexual dimorphism had been shown – or studied – in non-archosaurian sauropsids to our knowledge, many relevant examples are available in extant and sub-fossil archosaurs. A higher distal width in males than females was demonstrated in wild and captive *Alligator mississippiensis* using linear and geometric morphometrics (Farlow et al., 2005; Bonnan et al., 2008). Handley et al. (2016) demonstrated that femoral distal width of the more recently extinct flightless bird *Dromornis stirtoni* was also higher in males than females. To do so, they coupled morphometrics and multivariate statistics with the observation of medullary bone, a sex-specific tissue present in bones of egg-laying female in archosaurians (Dacke et al., 1993; Schweitzer et al., 2005, 2007; Canoville et al., 2019). The same kind of sexual dimorphism was observed in modern birds like California gulls (*Larus californicus*) (Schnell et al., 1985) and in the two extant species of ostriches (*Struthio c. camelus, S. c. molybdophanes*), but with reversed proportions between males and females (Elzanowski and Louchart, 2022). Furthermore, (Duggan et al., 2015) demonstrated that young male domestic ducks (*Anas platyrhynchos*) had more laterally curved femora than females, and that this sexually dimorphic feature disappeared along ontogeny. However, to our knowledge and aside Duggan et al. (2015), data about femoral obliquity is generally unavailable in most studies including sex determination in birds and other sauropsids. Therefore, because the femoral dimorphic features we highlighted in the Angeac-Charente ornithomimosaur herd were also demonstrated to vary between sexes in more or less closely related extant vertebrate clades, we infer it to be sexual.

We found no allometry along the first PC axis (Table S1), which indicates that the dimorphism we highlighted is not related to size. Ontogenetic allometry was often misinterpreted as sexual dimorphism in archosaurs, as demonstrated in the early dinosauriform *Asilisaurus kongwe*, the crocodylian *Alligator mississippiensis* and the bird *Rhea americana* (Griffin and Nesbitt, 2016; Hone and Mallon, 2017; Hedrick et al., 2021). Furthermore, this indicates no Sexual Size Dimorphism (SSD) in the Angeac-Charente ornithomimosaur. SSD is one of the most documented sexual dimorphism across all living organisms, whether it is biased toward females or males (Darwin, 1874; Fairbairn et al., 2007). There are many examples of observations and/or inferences of SSD and allometric relationships in extant and extinct dinosaurs (Larson, 1994; Bunce et al., 2003; Clarke, 2004; Székely et al., 2007; Remeš and Székely, 2010; Olson and Turvey, 2013; Handley et al., 2016; Manin et al., 2016; Fajemilehin, 2017). However, Elzanowski & Louchart (2022) demonstrated that female ostriches had more robust limb bones but smaller average body size than males. This decoupling between size and shape dimorphism is concordant with our results and emphasizes that sexual dimorphism is not necessarily reflected by body size nor allometry between limb segments. Thus, size-independent sexual dimorphism should be investigated further in extant archosaurs in order to improve inferences about sexual dimorphism in fossils, which are most often represented only by isolated bones.

We did not identify any other dimorphism in either the proximal part of the femur nor in complete tibia of the Angeac-Charente ornithomimosaurs (Fig. S2 & S3). However, sexual dimorphism was observed in the proximal ends of femora in extant ostriches (Charuta et al., 2007; Elzanowski and Louchart, 2022) and California gulls (Schnell et a l., 1985). In addition, the anteroposterior width of the femoral shaft was demonstrated to vary between sexes among savannah sparrows (*Passerculus sandwichensis*; Rising, 1987) and three species of steamer-ducks (*Tachyeres pteneres*, *T. leucocephalus*, *T. patachonicus*, (Livezey and Humphrey, 1984). Yet, and accordingly with our results, size-independent dimorphism in the avian tibiotarsus seems less common across the EPB. Indeed, to our knowledge, occurrences of shape dimorphism in the tibia was demonstrated only in California gulls (e.g., width of the shaft) (Schnell et al., 1985) and in ostriches [e.g., anteroposterior width of the distal epiphysis; only in Elzanowski & Louchart (2022) but not in Charuta et al. (2007)]. Furthermore, our observation that sexual dimorphism could be restricted to the femur in the Angeac-Charente ornithomimosaurs and modern archosaurs raises the question of the potential co-variation between the femur and the pelvis. Sexual dimorphism was observed in the ilium of several birds mentioned previously, such as ostriches, steamer-ducks, savannah sparrows, and California gulls (in the antitrochanter width, acetabular width and synsacrum width and length) (Livezey and Humphrey, 1984; Schnell et al., 1985; Rising, 1987; Charuta et al., 2007). All measurements were higher in male birds than in female birds except for the width of the ilium, which was higher in female ostriches when measured by Charuta et al. (2007), but not significantly different between sexes in Elzanowski & Louchart (2022). Additionally, female alligators had a deeper pelvic canal (i.e., distance between the ventral side of the first sacral vertebra and the ventral margin of the ischial symphysis) (Prieto-Marquez et al., 2007). The dimorphism was located preferably on the femur rather than on the tibia in the Angeac-Charente ornithomimosaur, which suggests that the pelvic area might as well be dimorphic, and that seems to be generally the case in some modern avian dinosaurs too (Livezey and Humphrey, 1984; Schnell et al., 1985; Rising, 1987; Farlow et al., 2005; Charuta et al., 2007; Prieto-Marquez et al., 2007; Bonnan et al., 2008; Duggan et al., 2015; Elzanowski and Louchart, 2022). Could the ability to carry egg restrict the location of sexual dimorphism closer to the hip region? Sexual dimorphism in the pelvic girdle, the proximal hindlimb and the morphological integration between the two in female extant archosaurs should be investigated further to answer this question.

Our results did not permit to confidently sex each morphotype. Most modern occurrences of femoral sexual dimorphism indicate a wider distal epiphysis among males than females, but Elzanowski & Louchart (2022) showed that the opposite was also true for modern and subfossils ostriches. Furthermore, our results indicated that femora with the narrowest distal epiphyses (females in most of modern occurrences) had a laterally deviated shaft. However, (Duggan et al., 2015) demonstrated that only juvenile male Pekin ducks had a laterally deviated shaft, which is not congruent with our results that the widest epiphyses were associated with a straighter morphotype. Paleohistological analyses could enable to verify sex assignment by assessing the presence of medullary bone, as some gravid females may have died during their egg-laying cycle at the time of the mass-mortality event recorded at Angeac-Charente. Indeed, medullary bone was recently demonstrated as probably the most reliable indicator of sex with an extensive distribution across the skeleton (Canoville et al., 2019). A paleohistological investigation could also confirm the ontogenetic homogeneity among our femoral sample, as recommended by Griffin & Nesbitt (2016), Hone & Mallon (2017) and Mallon (2017).

## Conclusion

Our results demonstrate that the femoral morphology among a large herd of coeval ornithomimosaurs is dimorphic. We identify bimodal distributions along size-independent features that were already reported to vary between sexes in modern archosaurs, and other tetrapods (*e.g*., the width of the distal epiphyses and the lateral deviation of the shaft). Therefore, we infer these features to indicate sexual dimorphism in the Angeac-Charente ornithomimosaurs according to the EPB approach. Our findings inform about the intraspecific variability in non-avian theropods and emphasize the need for description of size-independent dimorphism in modern and closely related taxa with a priori knowledge of the sex.

In the future, our results should be completed by paleohistological studies to 1) sex each morphotype and 2) identify the extent of ontogenetic variations within our sample. Additionally, we show that the sex-ratio of the Angeac-Charente ornithomimosaur is close to 1:1 and thus, likely Fisherian (Fisher, 1930). It was demonstrated that in extant archosaurs, Fisherian populations are only observed among clutches and hatchlings (Mayr, 1939; Clutton-Brock, 1986; Liker et al., 2013), and become generally biased toward females in sub-adult and adult populations, as demonstrated on crocodilians (Woodward and Murray, 1993; González et al., 2019) and ratites (Magige, 2012; Prokopenko et al., 2021). Therefore, paleohistological investigations could help characterize the variation of sex ratio along ontogeny in an extinct dinosaur population, and inform if it was truly Fisherian, unlike their extant relatives, or if it also experienced skewness along aging. More broadly, understanding how sex impacted the morphology of an extinct species could shed light on complex evolutionary mechanism such as trade-off between sexually dimorphic features, ecological adaptations and life-history traits.

## Material and Methods

### Sample and data acquisition

Several complete and fragmented femora and complete tibiae from the Angeac-Charente ornithomimosaur were discovered between 2010 and 2020 (Table 1). We removed 158 specimens that were too fragmented and altered by too much oxidized pyrite and trampling (femora: six complete, 37 proximal and 19 distal epiphyses; tibiae: four complete, 36 proximal and 56 distal epiphyses). We selected only fragmented femora that preserved: 1) the most proximal point of the fourth trochanter for proximal epiphyses; 2) the most proximal point of the anteromedial flange for distal epiphyses (Figure a). In total, we digitized 152 specimens (femora: 13 complete, 29 proximal and 21 distal epiphyses; tibiae: 21 complete, 30 proximal and 38 distal epiphyses) using the Artec EVA with Artec Studio Professional v. 12.1.1.12 (Artec 3D, Luxembourg, Luxembourg) and the NextEngine with Scan Studio Pro v. 2.0.2 (Next Engine inc., Santa Monica, United States) for a few specimens (Table S3). After re-examination of digitized specimens, we removed three complete femora, 14 proximal and eight distal epiphyses, and four complete tibiae that were distorted. We thus integrated 10 complete femora, 13 distal and 15 proximal femoral epiphyses, and 17 complete tibiae.

**Table 1.** Number of femora and tibiae from the Angeac-Charente ornithomimosaur discovered between 2010 and 2020. Minimum Number of Elements (MNE) and Minimum Number of Individuals (MNI) are given for each fragmented and complete femora.

|  | Femur | Tibia |
| --- | --- | --- |
| Left proximal (MNE) | 31 | 31 |
| Right proximal (MNE) | 35 | 35 |
| Left distal (MNE) | 18 | 48 |
| Right distal (MNE) | 22 | 46 |
| Left complete (MNE) | 8 | 13 |
| Right complete (MNE) | 11 | 12 |
| MNI | 46 | 61 |

### 3D geometric morphometrics

3D GMM is a well-established method for quantifying biological shape variations and has already enabled to identify sexual dimorphism in past studies (Kaliontzopoulou et al., 2007; Cavaignac et al., 2016). We followed a high-density morphometrics approach using a combination of single anatomical landmarks and sliding semilandmarks along curves and surfaces (Bookstein, 1997; Gunz et al., 2005). Indeed, most anatomical landmarks are usually concentrated on both ends of limb bones, hence why the use of sliding semilandmarks on surface was justified on the shaft (Gunz and Mitteroecker, 2013; Botton-Divet et al., 2016). We digitized 619 landmarks on complete femora (25 anatomical landmarks, 99 sliding semilandmarks on curves and 495 on surfaces), 479 on proximal (11 anatomical landmarks, 26 sliding semilandmarks on curves and 442 on surfaces) and distal epiphyses (10 anatomical landmarks, 45 sliding semilandmarks on curves and 424 on surfaces) and 725 on complete tibiae (23 anatomical landmarks, 219 sliding semilandmarks on curves and 483 on surfaces; see details in Figure S4; Table S4 & S5) using the IDAV Landmark software v. 3.0.0.6 (Wiley et al., 2005). We digitized anatomical landmarks and sliding semilandmarks along curves on each specimen and sliding semi-landmarks along surfaces on one specimen (ANG 10 90), referred to as “the template” hereafter (Cornette et al., 2013). We then automatically projected the sliding semilandmarks along surfaces of the template onto every other specimen following the spline relaxation of semilandmarks along curves using the function “placePatch” of the Morpho package v. 2.8 (Schlager, 2017). Then, we performed five iterations of another spline relaxation between landmark configurations of the template and the ones from every other specimen using the function “relaxLM” of Morpho. Finally, we performed a partial Procrustes fitting in order to compute a Procrustes consensus of every configuration and used it as a target for the two last iterations of spline relaxation using the function “slideLM” of Morpho. These three steps of spline relaxations ensured that every semilandmark position was geometrically homogeneous in all specimens (Gunz et al., 2005). Finally, we performed a Generalized Procrustes Analysis (GPA) using the function “gpagen” of the R package geomorph v. 3.3.1 (Adams and Otárola-Castillo, 2013) in order to align each femur in the Cartesian coordinate system by superimposing them based on their landmark configuration and to rule out the effect of size, location and orientation of the different landmark configurations (Gower, 1975; Rohlf and Slice, 1990; Zelditch et al., 2012).

### Statistical analyses and clustering

We performed a Principal Component Analysis (PCA) in order to reduce dimensionalities of the variation and isolate different components of shape variation (Gunz and Mitteroecker, 2013). The quantification of repeatability was performed by digitizing landmarks iteratively (n = 10) on three close specimens for complete femora and tibiae, which resulted in 30 configurations for each bone. We then computed a PCA for the two bones (30 configurations each), which showed that all 10 repetitions for each specimen were grouped together and isolated from those of the other specimens along the first two PC axes (Figure S5 & S6). This ensured that that biological variability was greater than the operator effect, which refers to the ability to reproduce accurately the same landmark configuration multiple times on the same specimen. As recommended by Mallon (Mallon, 2017), we performed mixture modelling analyses without a-priori knowledge about the number of groups in order to estimate how many morphological clusters would stand out in our dataset, if any, along each PC axis. Gaussians are well-suited functions to describe a biological population, especially when applied to a morphometric dataset (Baylac et al., 2003). We used the R package Mclust v. 5.4.7, which calculates the most-probable number of clusters in a dataset based on the detection of Gaussian distributions by maximum likelihood estimations (Scrucca et al., 2016). Bayesian Information Criteria (BIC; e.g., an approximation of Bayes factors for comparing likelihood) were used to choose which model, among the several ones available, fitted best with our dataset (i.e., the model with the highest BIC), while simultaneously estimating the number of Gaussian distributions (Fraley and Raftery, 2007). We computed 3D visualizations that highlighted which feature varied the most along each axis, and between clusters when dimorphism was identified. To do so, we first computed a 3D consensual mesh of all specimens of the sample by using the function “tps3d” from the R package Morpho v. 2.8 (Schlager, 2017) which performed a spline relaxation that minimized the bending energy of a Thin Plate Spline (TPS) between the template landmark configuration and a mean landmark configuration (obtained during the GPA). Then, the function used the resulting TPS deformation to warp the 3D mesh of the template onto the mean shape in order to compute a 3D consensual mesh (Bardua et al., 2019). Next, we calculated the mean coordinates of every specimen in each cluster along the PC axis identified as dimorphic by the mixture modelling analysis. Finally, we warped the mean shape, and its associated 3D mesh, onto the mean landmark configurations of each cluster by using the “shape.predictor” function of geomorphv. 3.3.1 (Adams and Otárola-Castillo, 2013) in order to visualize the 3D shape variation associated with the dimorphic PC axis. We studied the allometry within our sample [i.e., the size-related morphological variation (Klingenberg, 2016)], using Pearson’s correlation between each PC scores and the log-transformed centroid sizes using the R function “cor.test”.

## Acknowledgments

We warmly thank D. Augier, J.-F. Tournepiche (The museum of Angoulême) for access to specimens, A. Aumont, G. Baron (La Rochelle Museum) for access to specimens in temporary exhibit, F. Goussard (MNHN) for 3D digitizations with NextEngine and L. Rozada and J. Goedert (MNHN) for help with retrieving some specimens. We are indebted to the Audoin & Fils company for sharing their discovery with us and for providing us with logistical support. We would like to thank Mrs. Rodet, the Audoin family and the department of La Charente for donating all the fossil material discovered in Angeac-Charente to the museum of Angoulême. We acknowledge the contribution of professional and amateur paleontologists and the numerous students who participated in the Angeac-Charente excavations since 2010. We also thank B. Bed’hom. (MNHN) for very helpful discussions about sexual dimorphism in extant domestic birds. Finally, we thank C. Bader, A. Canoville, C. Etienne, J. Goedert, R. Lefebvre, K. Leverger, and C. Mallet for constructive discussions and recommendations about digitization, analyses and interpretation of data.

## Supplementary Figures and Tables

**Figure S1:**
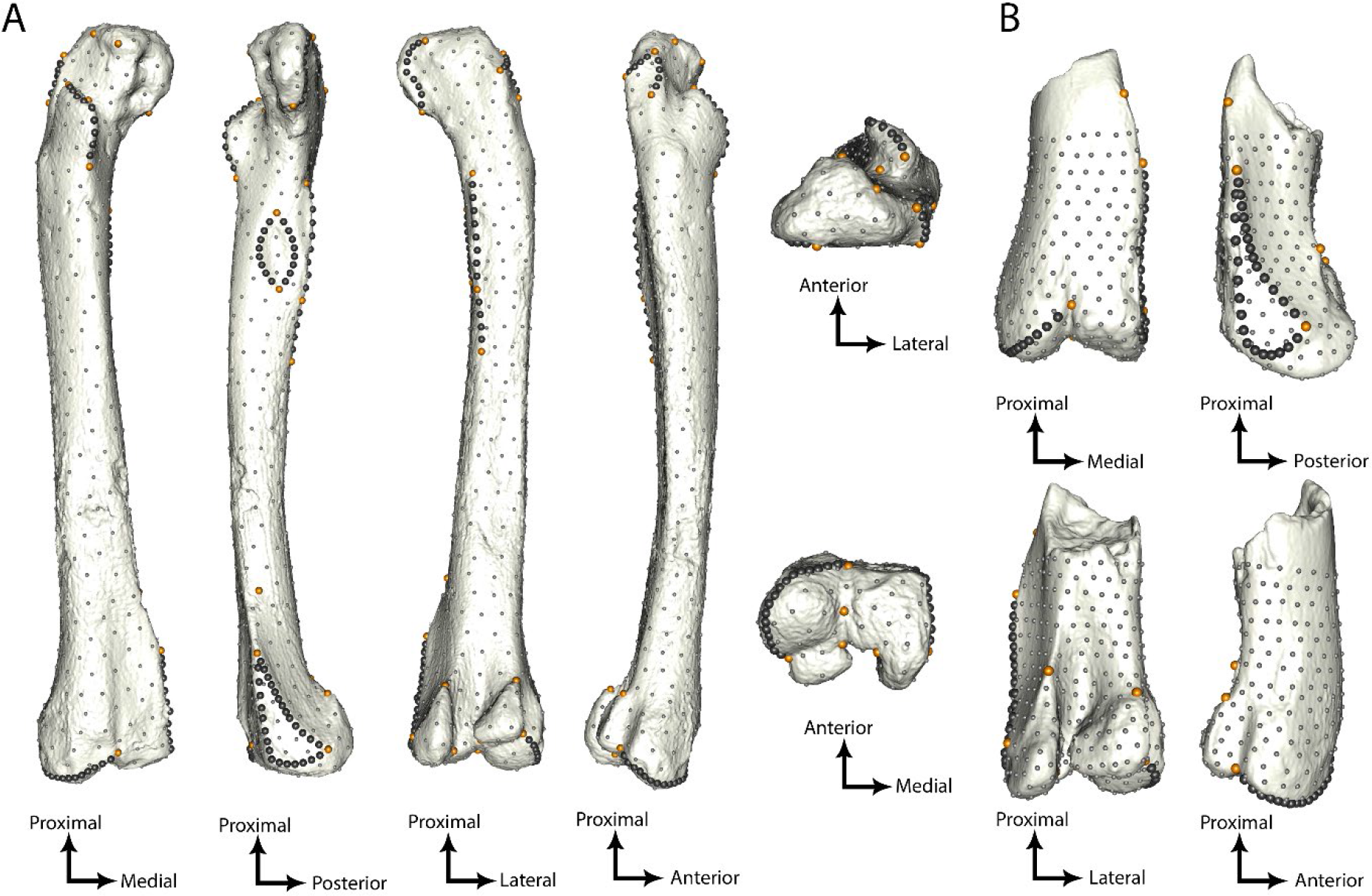
Template of A) right complete femur of ANG10 90 and B) mirrored left distal epiphysis of ANG14 3188 with anatomical landmarks (orange), sliding semilandmarks along curves (dark grey) and surfaces (light grey).

**Figure S2:**
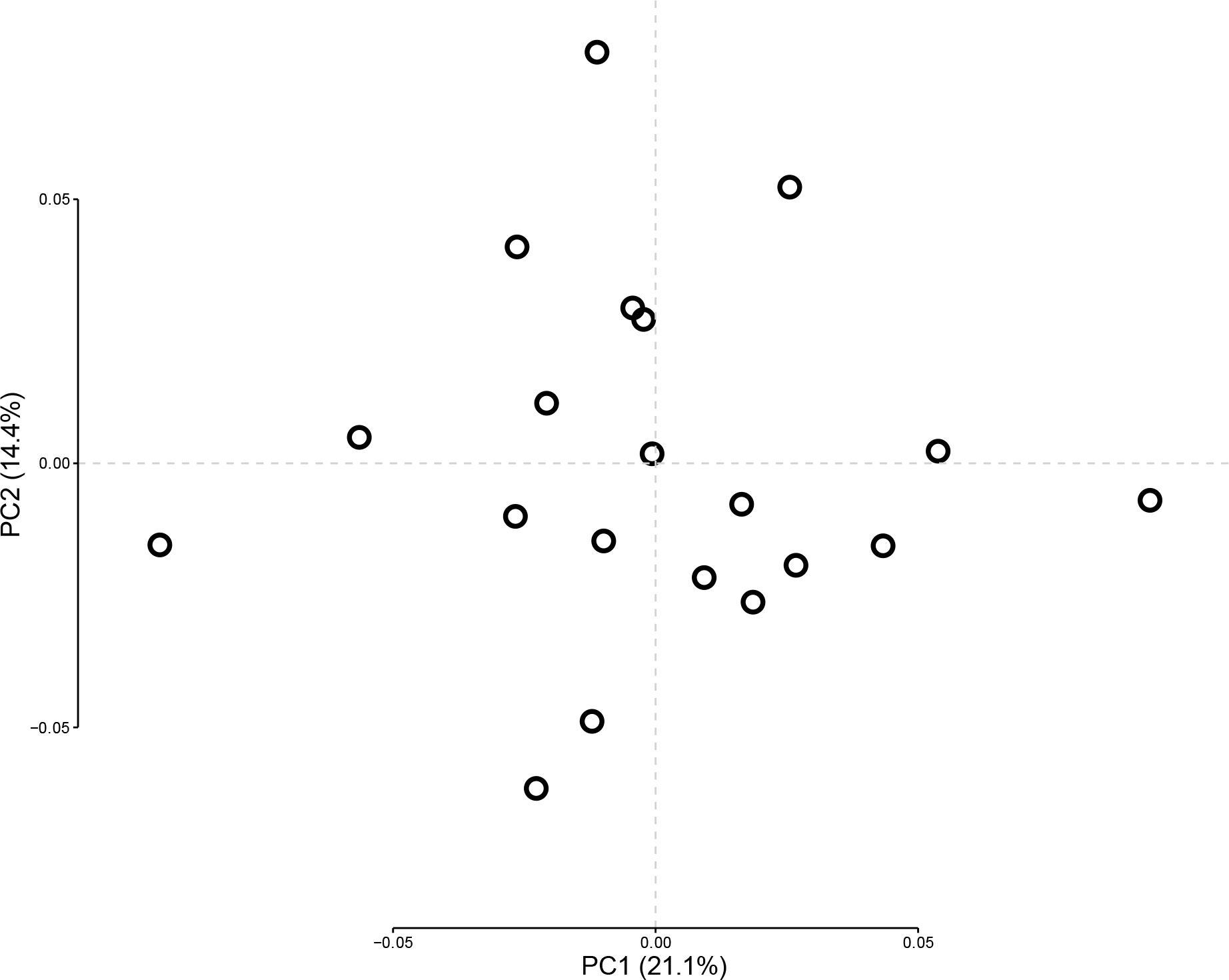
The two first axes of the PCA for proximal epiphyses of femora

**Figure S3:**
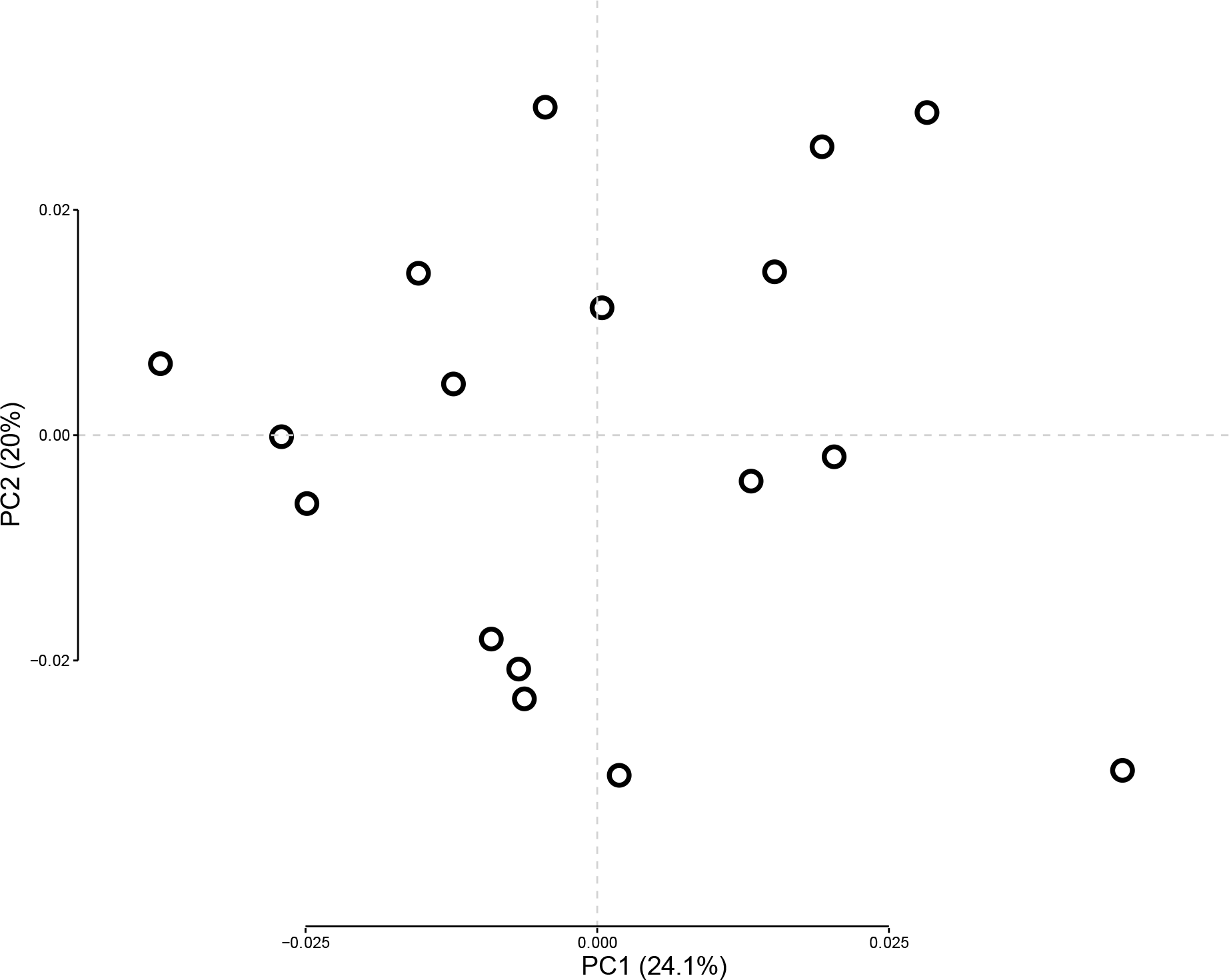
The two first axes of the PCA for complete tibiae.

**Figure S4:**
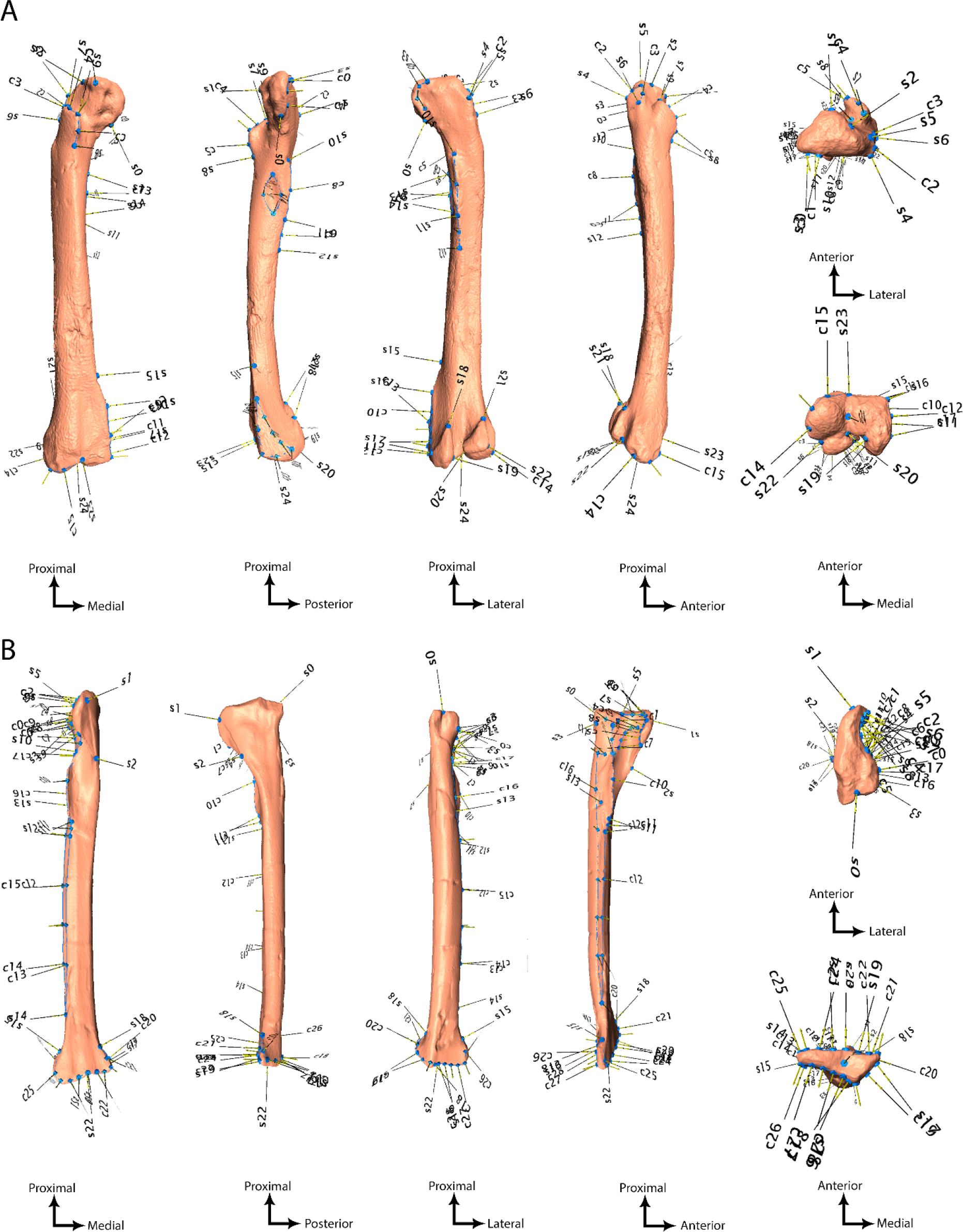
Landmark configuration on the templates A) femur; B) tibia, with numerotation following the scheme shown in Tables S4 & S5, Abbreviations: s, anatomical landmarks; c, sliding semilandmarks on curves.

**Figure S5:**
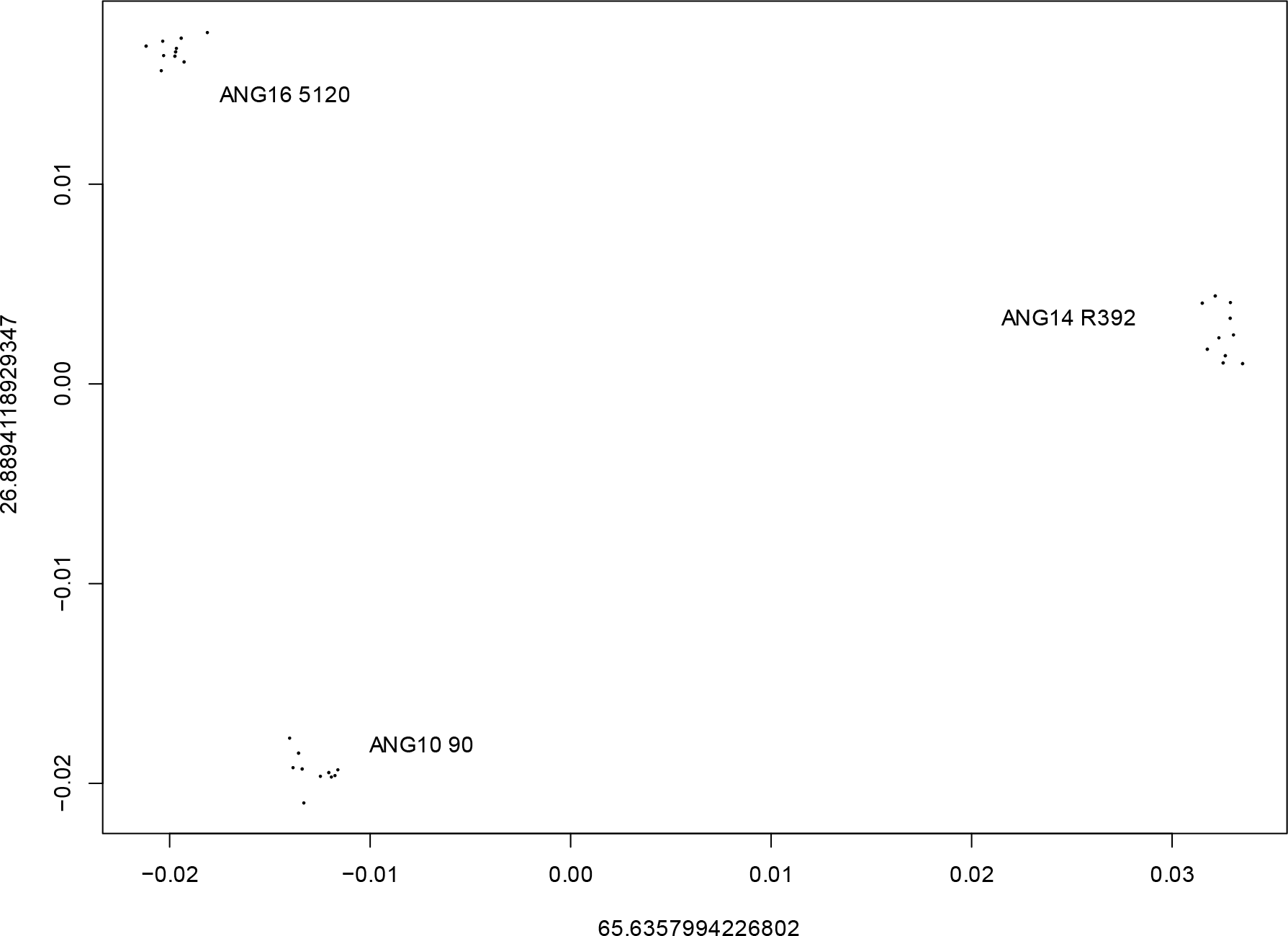
The two first axes of the PCA showing the quantification of the repeatability for the landmark configuration on femora.

**Figure S6:**
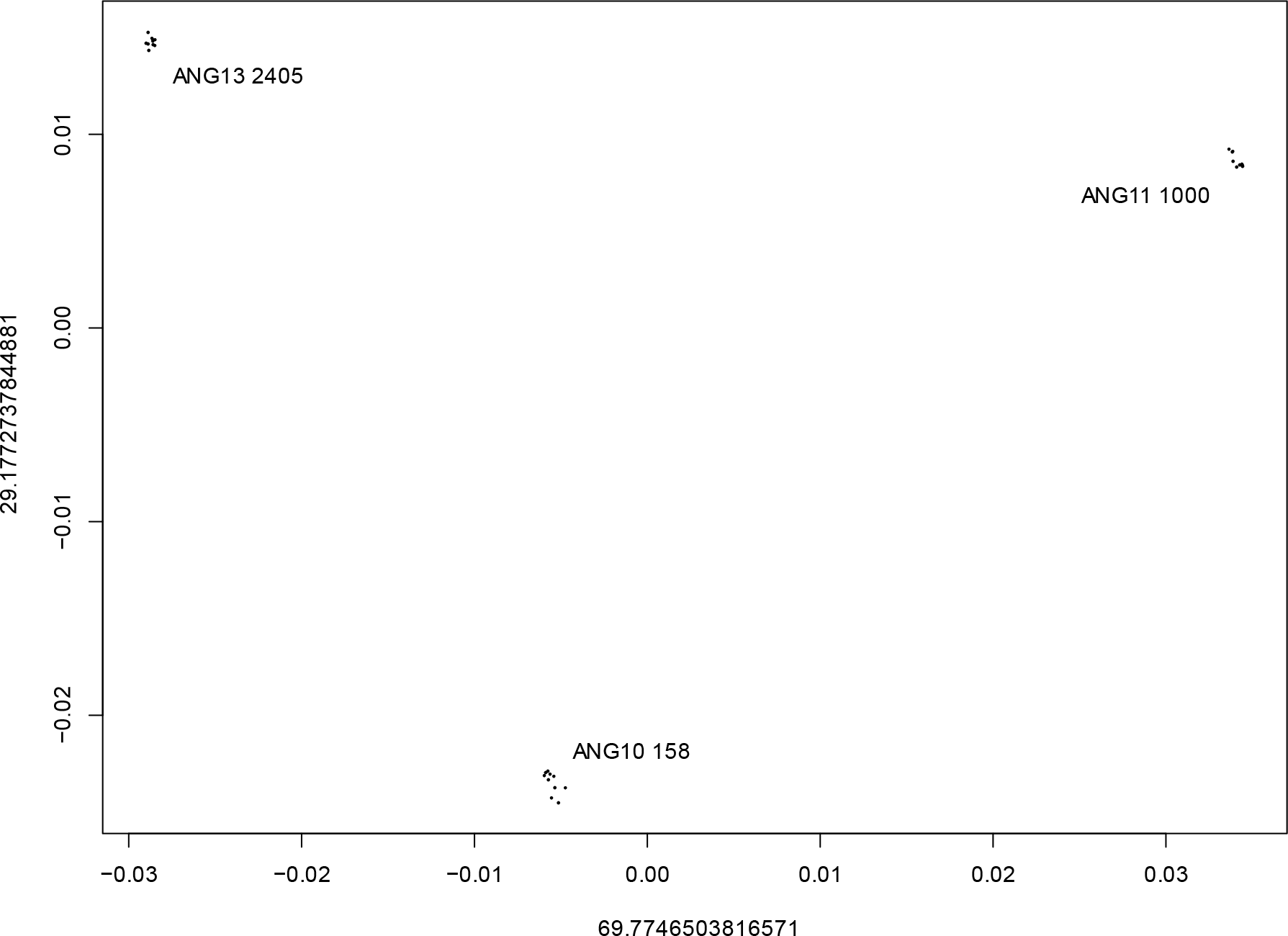
The two first axes of the PCA showing the quantification of the repeatability for the landmark configuration on tibiae.

**Table S1:** Statistical parameters used in this study for size-effect and cluster attribution

| Parameters | Complete femora | Distal epiphyses |
| --- | --- | --- |
| Log centroid size vs. PC1 scores | $r^2$: 0.12; $p$-value > 0.1 | $r^2$: 0.07 ; $p$-value > 0.1 |
| Model selected by the EM | univariate, equal variance | univariate, equal variance |
| Number of components | 2 | 2 |
| BIC | 46.54 | 48.47 |
| Log-likelihood | 27.87 | 30.12 |
| Mixing probabilities for each cluster | 0.61; 0.39 | 0.52; 0.48 |
| Highest uncertainty for cluster attribution/specimen | 0.0001 | 0.02 |

**Table S2:**
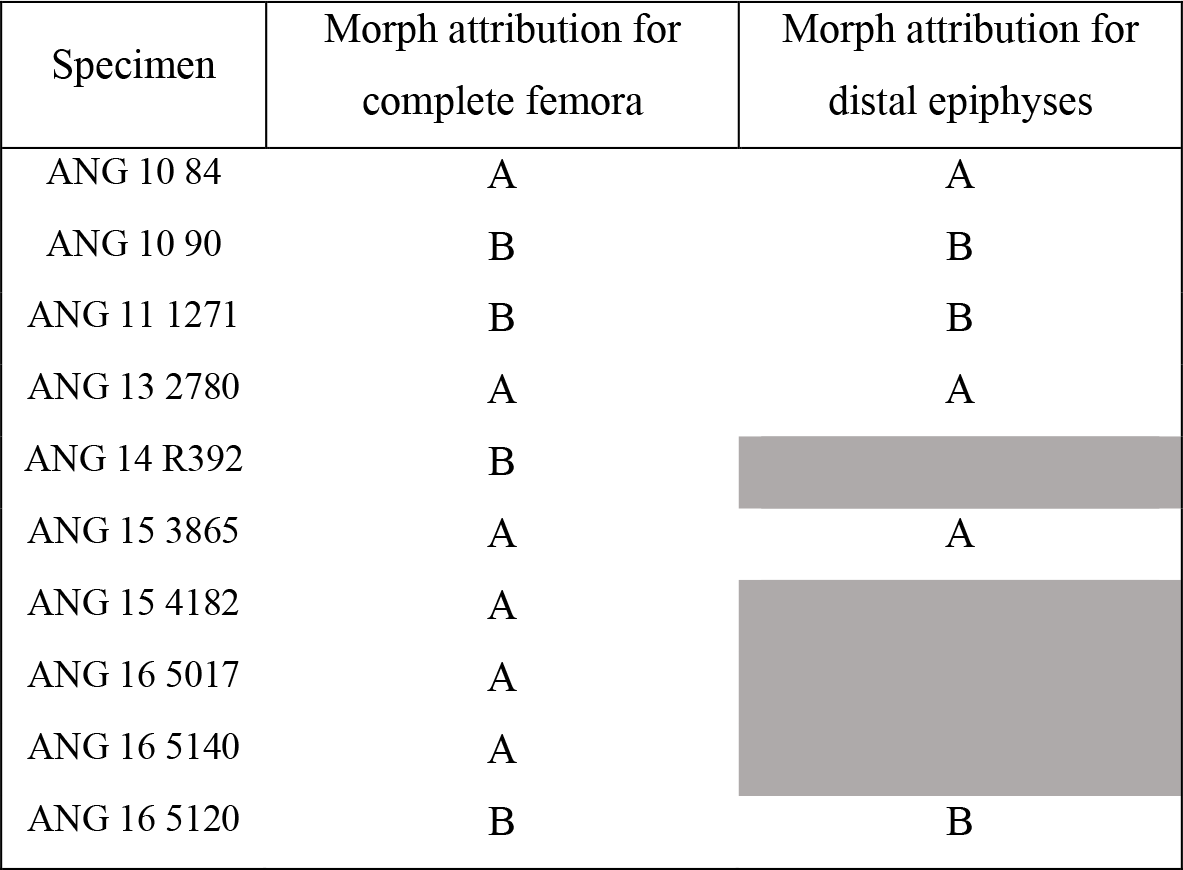
Cluster attribution for complete femora studied in analyses for both complete femora and distal epiphyses

**Table S3:** Specimens used in this study. * refers to specimens digitized with the NextEngine, other specimens were digitized using the Artec EVA. Abbreviations: Col. Nb., collection number; L, left; R, right

| Col. Nb. | Bone | Integrity | Side |
| --- | --- | --- | --- |
| ANG 10 43 | Femur | Proximal | L |
| ANG 10 53 | Femur | Proximal | R |
| ANG 10 84 | Femur | Complete | R |
| ANG 10 86 | Femur | Proximal | L |
| ANG 10 90 | Femur | Complete | L |
| ANG 10 171 | Femur | Distal | L |
| ANG 11 735 | Femur | Distal | R |
| ANG 11 811a | Femur | Proximal | R |
| ANG 11 811b | Femur | Distal | R |
| ANG 11 1107 | Femur | Distal | R |
| ANG 11 1209 | Femur | Proximal | L |
| ANG 11 1271 | Femur | Complete | R |
| ANG 12 1844 | Femur | Distal | L |
| ANG 13 2282 | Femur | Proximal | L |
| ANG 13 2381 | Femur | Proximal | R |
| ANG 13 2428 | Femur | Distal | L |
| ANG 13 2451 | Femur | Distal | R |
| ANG 13 2749 | Femur | Proximal | L |
| ANG 13 2757 | Femur | Proximal | R |
| ANG 13 2780 | Femur | Complete | L |
| ANG 13 2807 | Femur | Distal | R |
| ANG 14 R392 | Femur | Complete | R |
| ANG 14 3188 | Femur | Distal | L |
| ANG 14 3488 | Femur | Proximal | L |
| ANG 14 3516 | Femur | Proximal | R |
| ANG 14 3570 | Femur | Proximal | R |
| ANG 15 3865 | Femur | Complete | R |
| ANG 15 4182 | Femur | Complete | L |
| ANG 16 5017 | Femur | Complete | L |
| ANG 16 5106 | Femur | Proximal | R |
| ANG 16 5120 | Femur | Proximal | R |
| ANG 16 5140 | Femur | Complete | R |
| ANG 16 5077 | Femur | Distal | R |
| ANG 16 5120 | Femur | Complete | R |
| ANG 17 5704 | Femur | Proximal | L |
| ANG 17 5709 | Femur | Distal | L |
| ANG 19 6825* | Femur | Distal | L |
| ANG 20 7346* | Femur | Distal | R |
| ANG 10 158 | Tibia | Complete | L |
| ANG 10 1024.25 | Tibia | Complete | R |
| ANG 11 1000 | Tibia | Complete | R |
| ANG 12 1893 | Tibia | Complete | R |
| ANG 13 2405 | Tibia | Complete | L |
| ANG 13 2538 | Tibia | Complete | R |
| ANG 13 2588 | Tibia | Complete | L |
| ANG 13 2589 | Tibia | Complete | L |
| ANG 13 2599 | Tibia | Complete | L |
| ANG 13 2699 | Tibia | Complete | L |
| ANG 14 3031 | Tibia | Complete | L |
| ANG 14 3611 | Tibia | Complete | L |
| ANG 15 4038 | Tibia | Complete | L |
| ANG 15 4070 | Tibia | Complete | R |
| ANG 16 1349 | Tibia | Complete | R |
| ANG 16 5030a | Tibia | Complete | R |
| ANG 17 2207 | Tibia | Complete | L |

**Table S4:**
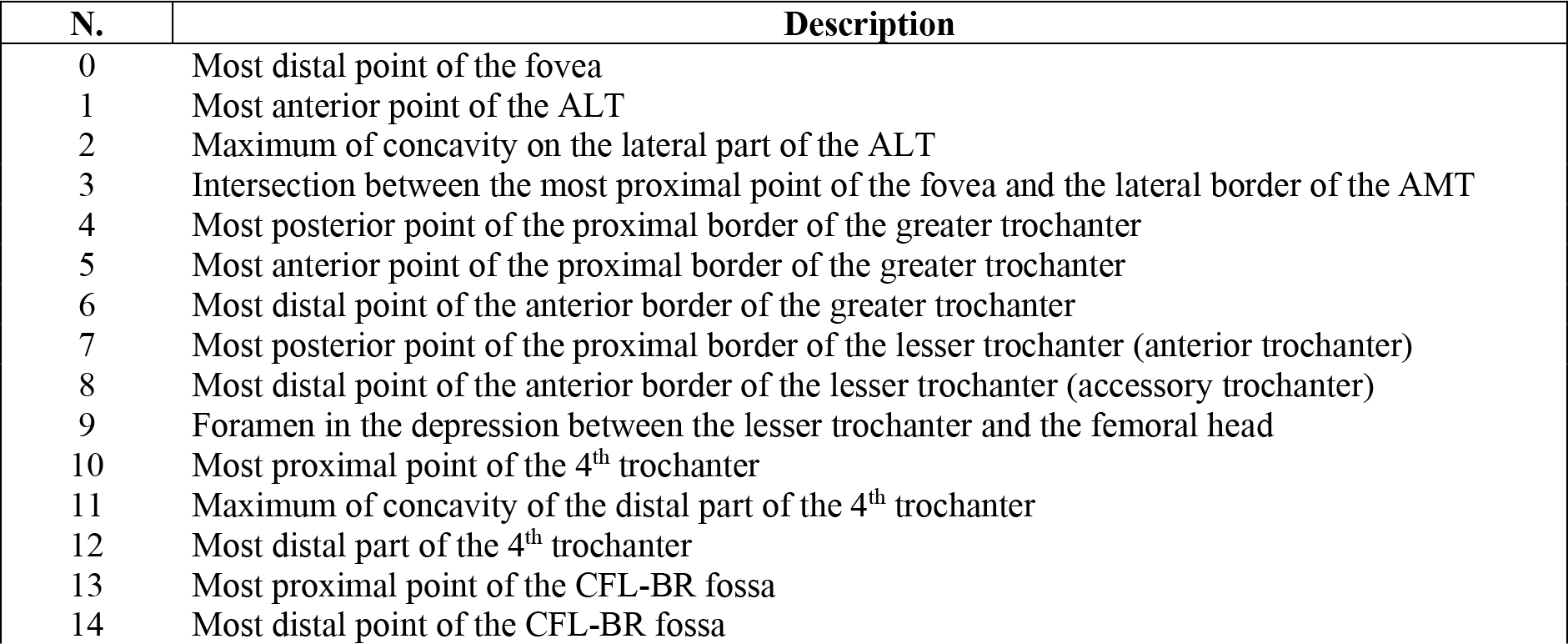
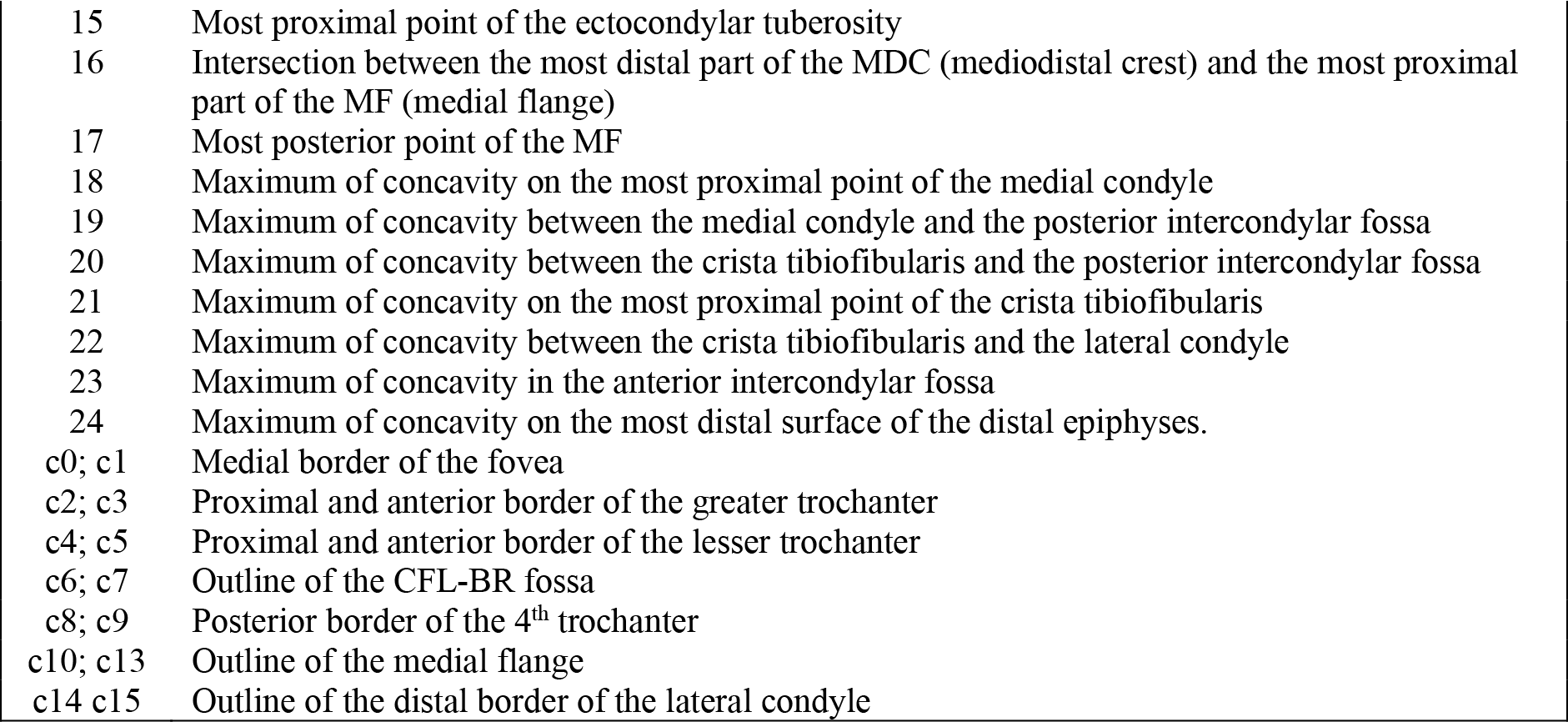
Landmark scheme of the femur according to the numerotation shown in Figure S4. Abbreviations: s, anatomical landmarks; c, sliding semilandmarks on curves.

**Table S5:** Landmark scheme of the tibia according to the numerotation shown in Figure S5. Abbreviations: s, anatomical landmarks; c, sliding semilandmarks on curves.

| N. | Description |
| --- | --- |
| 0 | Most proximal point of the maximum of concavity in the intercondylar groove on the tibial head |
| 1 | Most proximal point of the medial side of the cnemial crest |
| 2 | Maximum of concavity along the distal part of medial side of the cnemial crest |
| 3 | Most distal point of the lateral condyle |
| 4 | Most anterior point of the distal border of the lateral condyle |
| 5 | Most posterior point of the proximal border of the lateral side of the cnemial crest |
| 6 | Most anterior point of the proximal border of the lateral side of the cnemial crest |
| 7 | Most anterior point of the anterior border of the lateral side of the cnemial crest |
| 8 | Most posterior point of along the anterior border of the lateral side of the cnemial crest |
| 9 | Most distal point of the anterior border of the lateral side of the cnemial crest |
| 10 | Most proximal point of the fibular crest |
| 11 | Maximum of concavity of the distal part of the fibular crest |
| 12 | Most distal point of the fibular crest |
| 13 | Foramen on the posterior side of the fibular crest |
| 14 | Most distal point of the surface of contact with the fibula |
| 15 | Maximum of concavity on the proximal border of the lateral malleolus |
| 16 | Maximum of concavity along the lateral border of the posterior distal tuberosity |
| 17 | Most medial point of the medial malleolus |
| 18 | Maximum of concavity on the proximal border of the medial malleolus |
| 19 | Maximum of concavity along the medial border of the anterior distal tuberosity |
| 20 | Most anterior point of the anterior distal tuberosity |
| 21 | Maximum of concavity along the lateral border of the anterior distal tuberosity |
| 22 | Maximum of depression on the distal surfaces of the distal epiphysis |
| c0; c1 | Most distal border of the lateral side of the lateral condyle |
| c2; c9 | Outline of the fossa fibularis/insicular tibialis |
| c4; c5 | Proximal and anterior border of the lesser trochanter |
| c6; c7 | Outline of the CFL-BR fossa |
| c8; c9 | Posterior border of the 4<sup>th</sup> trochanter |
| c10; c11 | Outline of the fibular crest |
| c12; c17 | Outline of the surface of contact with the fibula |
| c18; c28 | Outline of the articular surface of the distal epiphysis |

## References

Adams, D.C., Otárola-Castillo, E., 2013. geomorph: an r package for the collection and analysis of geometric morphometric shape data. Methods in Ecology and Evolution 4, 393–399. https://doi.org/10.1111/2041-210X.12035

Allain, R., Vullo, R., Le Lœuff, J., Tournepiche, J.F., 2014. European ornithomimosaurs (Dinosauria, Theropoda): an undetected record. 105. https://doi.org/10.1344/105.000002083

Allain, R., Vullo, R., Rozada, L., Anquetin, J., Bourgeais, R., Goedert, J., Lasseron, M., Martin, J.E., Pérez-García, A., De Fabrègues, C.P., Royo-Torres, R., Augier, D., Bailly, G., Cazes, L., Despres, Y., Gailliègue, A., Gomez, B., Goussard, F., Lenglet, T., Vacant, R., Mazan, ., Tournepiche, J.-F., 2022. Vertebrate paleobiodiversity of the Early Cretaceous (Berriasian) Angeac-Charente Lagerstätte (southwestern France): implications for continental faunal turnover at the J/K boundary. Geodiversitas 44. https://doi.org/10.5252/geodiversitas2022v44a25

Alunni-Perret, V., Staccini, P., Quatrehomme, G., 2008. Sex determination from the distal part of the femur in a French contemporary population. Forensic Science International 175, 113–117. https://doi.org/10.1016/j.forsciint.2007.05.018

Bardua, C., Felice, R.N., Watanabe, A., Fabre, A.-C., Goswami, A., 2019. A Practical Guide to Sliding and Surface Semilandmarks in Morphometric Analyses. Integrative Organismal Biology 1, obz016. https://doi.org/10.1093/iob/obz016

Baylac, M., Villemant, C., Simbolotti, G., 2003. Combining geometric morphometrics with pattern recognition for the investigation of species complexes: Geometric morphometrics, pattern recognition and species complexes. Biological Journal of the Linnean Society 80, 89–98. https://doi.org/10.1046/j.1095-8312.2003.00221.x

Benton, M.J., Juul, L., Storrs, G.W., Galton, P.M., 2000. Anatomy and systematics of the prosauropod dinosaur *Thecodontosaurus antiquus* from the upper Triassic of southwest England. Journal of Vertebrate Paleontology 20, 77–108. https://doi.org/10.1671/0272-4634(2000)020[0077:AASOTP]2.0.CO;2

Bonnan, M.F., Farlow, J.O., Masters, S.L., 2008. Using linear and geometric morphometrics to detect intraspecific variability and sexual dimorphism in femoral shape in *Alligator mississippiensis* and its implications for sexing fossil archosaurs. Journal of Vertebrate Paleontology 28, 422–431. https://doi.org/10.1671/0272-4634(2008)28[422:ULAGMT]2.0.CO;2

Bookstein, F.L., 1997. Landmark methods for forms without landmarks: morphometrics of group differences in outline shape. Medical image analysis 1, 225–243.

Botton-Divet, L., Cornette, R., Fabre, A.-C., Herrel, A., Houssaye, A., 2016. Morphological Analysis of Long Bones in Semi-aquatic Mustelids and their Terrestrial Relatives. Integr. Comp. Biol. 56, 1298–1309. https://doi.org/10.1093/icb/icw124

Britt, B.B., Chure, D.J., Holtz Jr, T.R., Miles, C.A., Stadtman, K.L., 2000. A reanalysis of the phylogenetic affinities of Ceratosaurus (Theropoda, Dinosauria) based on new specimens from Utah, Colorado, and Wyoming. Journal of Vertebrate Paleontology 20.

Bunce, M., Worthy, T.H., Ford, T., Hoppitt, W., Willerslev, E., Drummond, A., Cooper, A., 2003. Extreme reversed sexual size dimorphism in the extinct New Zealand moa Dinornis. Nature 425, 172–175. https://doi.org/10.1038/nature01871

Canoville, A., Schweitzer, M.H., Zanno, L.E., 2019. Systemic distribution of medullary bone in the avian skeleton: ground truthing criteria for the identification of reproductive tissues in extinct Avemetatarsalia. BMC Evol Biol 19, 71. https://doi.org/10.1186/s12862-019-1402-7

Carrano, M.T., Sampson, S.D., Forster, C.A., 2002. The osteology of *Masiakasaurus knopfleri*, a small abelisauroid (Dinosauria: Theropoda) from the Late Cretaceous of Madagascar. Journal of Vertebrate Paleontology 22, 510–534. https://doi.org/10.1671/0272-4634(2002)022[0510:TOOMKA]2.0.CO;2

Cavaignac, E., Savall, F., Faruch, M., Reina, N., Chiron, P., Telmon, N., 2016. Geometric morphometric analysis reveals sexual dimorphism in the distal femur. Forensic Science International 259, 246.e1–246.e5. https://doi.org/10.1016/j.forsciint.2015.12.010

Chapman, R.E., Weishampel, D.B., Hunt, G., Rasskin-Gutman, D., 1997. Sexual dimorphism in dinosaurs, in: Dinofest International. pp. 83–93.

Charuta, A., Dzierzęcka, G., Reymond, J., Mańvkowska-Pliszka, H., 2007. Morfologia i morfometria obręczy oraz części wolnej kończyny miednicznej strusia. Medycyna Weterynaryjna 63, 1090–1094.

Clarke, J.A., 2013. Feathers before flight. Science 340, 690–692.

Clarke, J.A., 2004. Morphology, phylogenetic taxonomy, and systematics of Ichthyornis and Apatornis (Avialae: Ornithurae). amnb 2004, 1–179. https://doi.org/10.1206/0003-0090(2004)286<0001:MPTASO>2.0.CO;2

Clutton-Brock, T.H., 1986. Sex ratio variation in birds. Ibis 128, 317–329.

Colbert, E.H., 1990. Variation in Coelophysis bauri. En: K. Carpenter & PJ Currie (Eds.), Dinosaur systematics: approaches and perpectives.

Cornette, R., Baylac, M., Souter, T., Herrel, A., 2013. Does shape co-variation between the skull and the mandible have functional consequences? A 3D approach for a 3D problem. J. Anat. 223, 329–336. https://doi.org/10.1111/joa.12086

Cox, R.M., Butler, M.A., John-Alder, H.B., 2007. The evolution of sexual size dimorphism in reptiles, in: Fairbairn, D.J., Blanckenhorn, W.U., Székely, T. (Eds.), Sex, Size and Gender Roles: Evolutionary Studies of Sexual Size Dimorphism. Oxford University Press, pp. 38–49.

Dacke, C.G., Arkle, S., Cook, D.J., Wormstone, I.M., Jones, S., Zaidi, M., Bascal, Z.A., 1993. Medullary bone and avian calcium regulation 26.

Darwin, C.R., 1874. The descent of man, and selection in relation to sex. 2nd edn. John Murray, London.

Dodson, P., 1976. Taxonomic implications of relative growth in lambeosaurinae hadrosaurs. Systematic Zoology 24, 37–54.

Dong, Z., 1997. Mixture Analysis and its Preliminary Application in Archaeology. Journal of Archaeological Science 24, 141–162. https://doi.org/10.1006/jasc.1996.0100

Duggan, B.M., Hocking, P.M., Schwarz, T., Clements, D.N., 2015. Differences in hindlimb morphology of ducks and chickens: effects of domestication and selection. Genet Sel Evol 47, 88. https://doi.org/10.1186/s12711-015-0166-9

Dunn, P.O., Whittingham, L.A., Pitcher, T.E., 2001. Mating systems, sperm competition, and the evolution of sexual dimorphism in birds. Evolution 55, 161–175.

Elzanowski, A., Louchart, A., 2022. Metric variation in the postcranial skeleton of ostriches, *Struthio* (Aves: Palaeognathae), with new data on extinct subspecies. Zoological Journal of the Linnean Society 195, 88–105. https://doi.org/10.1093/zoolinnean/zlab049

Fabre, A.-C., Cornette, R., Huyghe, K., Andrade, D.V., Herrel, A., 2014. Linear versus geometric morphometric approaches for the analysis of head shape dimorphism in lizards: Head Shape Dimorphism in *Tupinambis*. Journal of Morphology 275, 1016–1026. https://doi.org/10.1002/jmor.20278

Fairbairn, D.J., Blanckenhorn, W.U., Székely, T. (Eds.), 2007. Sex, size, and gender roles: evolutionary studies of sexual size dimorphism, Oxford biology. Oxford University Press, Oxford ; New York.

Fajemilehin, S.O.K., 2017. Discriminant analysis of sexual dimorphism in zoometrical characters of normal feathered Yoruba ecotype adult local chicken in the Tropical Forest Zone of Nigeria. JASVM 2, 139–144. https://doi.org/10.31248/JASVM2017.060

Farlow, J.O., Hurlburt, G.R., Elsey, R.M., Britton, A.R.C., Langston, W., 2005. Femoral dimensions and body size of *Alligator mississippiensis*: estimating the size of extinct mesoeucrocodylians. Journal of Vertebrate Paleontology 25, 354–369. https://doi.org/10.1671/0272-4634(2005)025[0354:FDABSO]2.0.CO;2

Fisher, R.A., 1930. The Genetical Theory of Natural Selection. Oxford, UK: Clarendon Press.

Fitch, H.S., 1981. Sexual size differences in reptiles. Miscellaneous publications of the Museum of Natural History University of Kansas 70, 1–72.

Fraley, C., Raftery, A.E., 2007. Bayesian regularization for normal mixture estimation and model-based clustering. Journal of classification 24, 155–181.

Gaikwad, K.R., Nikam, V.R., 2014. Sexual dimorphism in femur. IOSR Journal of Dental and Medical Sciences 13, 4–9.

Godfrey, L.R., Lyon, S.K., Sutherland, M.R., 1993. Sexual dimorphism in large-bodied primates: the case of the subfossil lemurs. American Journal of Physical Anthropology 90, 315–334.

González, E.J., Martínez-López, M., Morales-Garduza, M.A., García-Morales, R., Charruau, P., Gallardo-Cruz, J.A., 2019. The sex-determination pattern in crocodilians: A systematic review of three decades of research. Journal of Animal Ecology 88, 1417–1427.

Gower, J.C., 1975. Generalized procrustes analysis. Psychometrika 40, 33–51. https://doi.org/10.1007/bf02291478

Griffin, C.T., Nesbitt, S.J., 2016. The femoral ontogeny and long bone histology of the Middle Triassic (?late Anisian) dinosauriform *Asilisaurus kongwe* and implications for the growth of early dinosaurs. Journal of Vertebrate Paleontology 36, e1111224. https://doi.org/10.1080/02724634.2016.1111224

Gunz, P., Mitteroecker, P., 2013. SEMILANDMARKS: A METHOD FOR QUANTIFYING CURVES AND SURFACES. Hystrix, the Italian Journal of Mammalogy 24. https://doi.org/10.4404/hystrix-24.1-6292

Gunz, P., Mitteroecker, P., Bookstein, F.L., 2005. Semilandmarks in Three Dimensions, in: Slice, D.E. (Ed.), Modern Morphometrics in Physical Anthropology. Kluwer Academic Publishers-Plenum Publishers, New York, pp. 73–98. https://doi.org/10.1007/0-387-27614-9_3

Handley, W.D., Chinsamy, A., Yates, A.M., Worthy, T.H., 2016. Sexual dimorphism in the late Miocene mihirung *Dromornis stirtoni* (Aves: Dromornithidae) from the Alcoota Local Fauna of central Australia. Journal of Vertebrate Paleontology 36, e1180298. https://doi.org/10.1080/02724634.2016.1180298

Hedrick, B.P., Schachner, E.R., Dodson, P., 2021. Alligator appendicular architecture across an ontogenetic niche shift. Anat Rec ar.24717. https://doi.org/10.1002/ar.24717

Hone, D.W.E., Mallon, J.C., 2017. Protracted growth impedes the detection of sexual dimorphism in non-avian dinosaurs. Palaeontology 60, 535–545. https://doi.org/10.1111/pala.12298

Hone, D.W.E., Mallon, J.C., Hennessey, P., Witmer, L.M., 2020. Ontogeny of a sexually selected structure in an extant archosaur *Gavialis gangeticus* (Pseudosuchia: Crocodylia) with implications for sexual dimorphism in dinosaurs. PeerJ 8, e9134. https://doi.org/10.7717/peerj.9134

Hunt, K.D., Dunevant, S.E., Yohler, R.M., Carlson, K.J., 2021. Femoral Bicondylar Angles among Dry-Habitat Chimpanzees (Pan troglodytes schweinfurthii) Resemble Those of Humans: Implications for Knee Function, Australopith Sexual Dimorphism, and the Evolution of Bipedalism. Journal of Anthropological Research 77, 303–337.

Kaliontzopoulou, A., Carretero, M.A., Llorente, G.A., 2007. Multivariate and geometric morphometrics in the analysis of sexual dimorphism variation inPodarcis lizards. J. Morphol. 268, 152–165. https://doi.org/10.1002/jmor.10494

Klingenberg, C.P., 2016. Size, shape, and form: concepts of allometry in geometric morphometrics. Dev Genes Evol 226, 113–137. https://doi.org/10.1007/s00427-016-0539-2

Knell, R.J., Naish, D., Tomkins, J.L., Hone, D.W.E., 2013. Sexual selection in prehistoric animals: detection and implications. Trends in Ecology & Evolution 28, 38–47. https://doi.org/10.1016/j.tree.2012.07.015

Knell, R.J., Sampson, S., 2011. Bizarre structures in dinosaurs: species recognition or sexual selection? A response to Padian and Horner. Journal of Zoology 283, 18–22. https://doi.org/10.1111/j.1469-7998.2010.00758.x

Larson, P.L., 1994. Tyrannosaurus sex. The Paleontological Society Special Publications 7, 139–156.

Liker, A., Freckleton, R.P., Székely, T., 2013. The evolution of sex roles in birds is related to adult sex ratio. Nature communications 4, 1–6.

Livezey, B.C., Humphrey, P.S., 1984. Sexual Dimorphism in Continental Steamer-Ducks. The Condor 86, 368–377. https://doi.org/10.2307/1366809

Magige, F.J., 2012. Spatial-temporal variation in sex ratio and group size of ostriches (Struthio namelus) in the Serengeti National Park and environs in Northern Tanzania. Tanzania Journal of Science 38, 15–23.

Mallon, J.C., 2017. Recognizing sexual dimorphism in the fossil record: lessons from nonavian dinosaurs. Paleobiology 43, 495–507. https://doi.org/10.1017/pab.2016.51

Manin, A., Cornette, R., Lefèvre, C., 2016. Sexual dimorphism among Mesoamerican turkeys: A key for understanding past husbandry. Journal of Archaeological Science: Reports 10, 526–533. https://doi.org/10.1016/j.jasrep.2016.05.066

Mayr, E., 1939. The sex ratio in wild birds. The American Naturalist 73, 156–179.

Morris, J.S., Brandt, E.K., 2014. Specialization for aggression in sexually dimorphic skeletal morphology in grey wolves (*Canis lupus*). J. Anat. 225, 1–11. https://doi.org/10.1111/joa.12191

Morris, J.S., Carrier, D.R., 2016. Sexual selection on skeletal shape in Carnivora. Evolution 70, 767–780.

Olson, V.A., Turvey, S.T., 2013. The evolution of sexual dimorphism in New Zealand giant moa (*Dinornis*) and other ratites. Proc. R. Soc. B. 280, 20130401. https://doi.org/10.1098/rspb.2013.0401

Owens, I.P.F., Hartley, I.R., 1998. Sexual dimorphism in birds: why are there so many different forms of dimorphism? Proc. R. Soc. Lond. B 265, 397–407. https://doi.org/10.1098/rspb.1998.0308

Padian, K., Horner, J.R., 2011. The evolution of ‘bizarre structures’ in dinosaurs: biomechanics, sexual selection, social selection or species recognition? Journal of Zoology 283, 3–17. https://doi.org/10.1111/j.1469-7998.2010.00719.x

Parsons, F.G., 1914. The characters of the English thigh-bone. Journal of Anatomy and physiology 48, 238.

Piechowski, R., Tałanda, M., Dzik, J., 2014. Skeletal variation and ontogeny of the Late Triassic Dinosauriform *Silesaurus opolensis*. Journal of Vertebrate Paleontology 34, 1383–1393. https://doi.org/10.1080/02724634.2014.873045

Prieto-Marquez, A., Gignac, P.M., Joshi, S., 2007. Neontological evaluation of pelvic skeletal attributes purported to reflect sex in extinct non-avian archosaurs. Journal of Vertebrate Paleontology 27, 603–609. https://doi.org/10.1671/0272-4634(2007)27[603:NEOPSA]2.0.CO;2

Prokopenko, N., Melnyk, V., Bazyvoliak, S., 2021. Biological features of egg productivity of black African ostriches under a semi-intensive keeping. Ukrainian Journal of Ecology 33–36.

Raath, M.A., Carpenter, K., Currie, P.J., 1990. Morphological variation in small theropods and its meaning in systematics: evidence from Syntarsus. Carpenter and Currie 91–105.

Remeš, V., Székely, T., 2010. Domestic chickens defy Rensch’s rule: sexual size dimorphism in chicken breeds: Sexual size dimorphism in chicken breeds. Journal of Evolutionary Biology 23, 2754–2759. https://doi.org/10.1111/j.1420-9101.2010.02126.x

Rising, J.D., 1987. GEOGRAPHIC VARIATION OF SEXUAL DIMORPHISM IN SIZE OF SAVANNAH SPARROWS (*PASSERCULUS SANDWICHENSIS*): A TEST OF HYPOTHESES. Evolution 41, 514–524. https://doi.org/10.1111/j.1558-5646.1987.tb05822.x

Rohlf, F.J., Slice, D., 1990. Extensions of the Procrustes Method for the Optimal Superimposition of Landmarks. Systematic Biology 39, 40–59. https://doi.org/10.2307/2992207

Rozada, L., Allain, R., Vullo, R., Goedert, J., Augier, D., Jean, A., Marchal, J., Peyre de Fabrègues, C., Qvarnström, M., Royo-Torres, R., 2021. A Lower Cretaceous *Lagerstätte* from France: a taphonomic overview of the Angeac-Charente vertebrate assemblage. Lethaia 54, 141–165. https://doi.org/10.1111/let.12394

Rozada, L., Allain, R., Vullo, R., Leprince, A., Tournepiche, J.F., 2014. Taphonomy of the ornithomimosaur dinosaur herd from the Early Cretaceous lignitic bone bed of Angeac-Charente (France), in: 74th Annual Meeting of the Society of Vertebrate Paleontology.

Saitta, E.T., 2015. Evidence for Sexual Dimorphism in the Plated Dinosaur Stegosaurus mjosi (Ornithischia, Stegosauria) from the Morrison Formation (Upper Jurassic) of Western USA. PLoS ONE 10, e0123503. https://doi.org/10.1371/journal.pone.0123503

Saitta, E.T., Stockdale, M.T., Longrich, N.R., Bonhomme, V., Benton, M.J., Cuthill, I.C., Makovicky, P.J., 2020. An effect size statistical framework for investigating sexual dimorphism in non-avian dinosaurs and other extinct taxa. Biological Journal of the Linnean Society 131, 231–273. https://doi.org/10.1093/biolinnean/blaa105

Schlager, S., 2017. Morpho and Rvcg – Shape Analysis in R, in: Statistical Shape and Deformation Analysis. Elsevier, pp. 217–256. https://doi.org/10.1016/B978-0-12-810493-4.00011-0

Schnell, G.D., Worthen, G.L., Douglas, M.E., 1985. Morphometric Assessment of Sexual Dimorphism in Skeletal Elements of California Gulls. The Condor 87, 484–493. https://doi.org/10.2307/1367944

Schweitzer, M.H., Elsey, R.M., Dacke, C.G., Horner, J.R., Lamm, E.-T., 2007. Do egg-laying crocodilian (*Alligator mississippiensis*) archosaurs form medullary bone? Bone 40, 1152–1158. https://doi.org/10.1016/j.bone.2006.10.029

Schweitzer, M.H., Wittmeyer, J.L., Horner, J.R., 2005. Gender-Specific Reproductive Tissue in Ratites and *Tyrannosaurus rex*. Science 308, 1456–1460. https://doi.org/10.1126/science.1112158

Scrucca, L., Fop, M., Murphy, T.B., Raftery, A.E., 2016. mclust 5: clustering, classification and density estimation using Gaussian finite mixture models. The R journal 8, 289.

Székely, Tamas, Lislevand, T., Figuerola, J., 2007. Sexual size dimorphism in birds, in: Fairbairn, D.J., Blanckenhorn, W.U., Székely, Tamás (Eds.), Sex, Size and Gender Roles: Evolutionary Studies of Sexual Size Dimorphism. Oxford University Press, pp. 27–37.

Tardieu, C., Glard, Y., Garron, E., Boulay, C., Jouve, J.-L., Dutour, O., Boëtsch, G., Bollini, G., 2006. Relationship between formation of the femoral bicondylar angle and trochlear shape: independence of diaphyseal and epiphyseal growth. American Journal of Physical Anthropology: The Official Publication of the American Association of Physical Anthropologists 130, 491–500.

Wiley, D.F., Amenta, N., Alcantara, D.A., Ghosh, D., Kil, Y.J., Delson, E., Harcourt-Smith, W., Rohlf, F.J., St John, K., Hamann, B., 2005. Evolutionary morphing. IEEE.

Witmer, L.M., Thomason, J.J., 1995. The extant phylogenetic bracket and the importance of reconstructing soft tissues in fossils. Functional morphology in vertebrate paleontology 1, 19–33.

Woodward, D.E., Murray, J.D., 1993. On the effect of temperature-dependent sex determination on sex ratio and survivorship in crocodilians. Proceedings of the Royal Society of London. Series B: Biological Sciences 252, 149–155.

Zelditch, M.L., Swiderski, D.L., Sheets, H.D., Fink, W.L. (Eds.), 2012. Geometric morphometrics for biologists: a primer. Elsevier Academic Press, London, Waltham, San Diego:

